# Echoes of self: Understanding acoustic structure and informational content in common marmoset (*Callithrix jacchus*) phee sequences

**DOI:** 10.1101/2024.04.14.589400

**Authors:** Kristin Meshinska, Judith M Burkart, Matthew BV Bell, Kaja Wierucka

**Affiliations:** Institute of Evolutionary Biology, School of Biological Sciences, University of Edinburgh, Edinburgh Edinburgh, United Kingdom; Department of Evolutionary Anthropology, University of Zurich, Winterthurerstrasse 190, 8057 Zurich, Switzerland; Center for the Interdisciplinary Study of Language Evolution, University of Zurich, Affolternstrasse 56, 8050 Zürich, Switzerland; German Primate Center - Leibniz Institute for Primate Research, Behavioural Ecology and Sociobiology Unit, Kellnerweg 4, 37077 Göttingen, Germany

## Abstract

Communication in social animals relies heavily on acoustic cues, yet many species possess limited vocal repertoires. To compensate, animals often produce vocalizations in sequences, potentially enhancing the diversity of transmitted information. However, the significance of repeated elements within these sequences remains poorly understood. Here, we investigated the spectro-temporal structure of elements within common marmoset (*Callithrix jacchus*) phees, a long-distance contact call, often produced in sequences. Employing machine learning techniques (random forests) and linear mixed effects models, we explored how elements varied based on their position within sequences and assessed their ability to encode identity and sex information. Additionally, we examined similarities between elements occupying the same position in different sequences. Our results reveal structural differences both within and between sequences, with variations observed in phees at different positions within the same sequence, yet similarities found between first elements of distinct sequences. Notably, all phees encoded caller identity and sex information, with varying accuracy across positions within sequences, indicating a higher encoding of sex information at the beginning of the sequence and a greater emphasis on identity in subsequent elements. These findings suggest that repeated sequences may be functionally diverse structures, enriching the complexity of animal communication systems.

## Introduction

During its lifetime an individual encounters and interacts with a variety of individuals in different social contexts. Effective communication between these parties is essential to mediate social behaviors, such as group coordination, reproduction, parental care, and competition (Clutton-Brock, 2016). Acoustic signals and cues can encode both dynamic (e.g., short-term fluctuations in physiological and psychological states) and static information, that is stable over long periods of time (e.g., identity, sex, age; Charlton et al., 2020). This information can be useful in many social contexts and has been studied mainly as a medium for parent-offspring recognition, mate selection, sexual competition, kin discrimination, and territoriality (Taylor et al., 2016).

Individual identity is amongst the most important pieces of information conveyed in the social contexts outlined above, as it allows receivers to weigh the risks and benefits of interacting with a given individual. It is especially useful when these interactions are likely to repeat (Tibbetts and Dale, 2007). In vocal communication, identity signatures can arise through anatomical differences between individuals, such as differences in the morphology of the vocal apparatus (Charlton et al., 2020). These differences are then manifested through variation in either one or a combination of spectral-temporal features, such as fundamental and formant frequencies (Taylor et al., 2016).

The vocalizations of many vertebrates provide information about the caller’s sex (Riebel et al., 2019; Serrano et al., 2020). Generally, sex differences in acoustic cues are advantageous when an individual is trying to attract a mate (Charlton et al., 2007; Martin et al., 2017; Semple and McComb, 2000). They also mediate potentially aggressive social interactions such as competition for resources (e.g., nesting sites) and breeding positions, or mate guarding (Riebel et al., 2019; Rubow et al., 2017; Serrano et al., 2020). Sex differences in the acoustic cues of dimorphic species are commonly explained through body size and anatomical differences between the sexes which manifest as vocal frequency variation (Charlton et al., 2020; Ey et al., 2007; Rendall et al., 2004). Following this theory, we might not expect to find sex differences in the vocalizations of monomorphic species. However this does not seem to be the case, as suggested by research on species such as cottontop tamarins, *Saguinus oedipus* (Miller et al., 2004), indri lemurs, *Indri indri* (Giacoma et al., 2010), dwarf mongooses, *Helogale parvula* (Rubow et al., 2016), and rock hyraxes, *Procavia capensis* (Frydman et al., 2023) where sex differences may occur due to differences in hormonal profiles, morphological differences of the vocal apparatus or due to variation in calling patterns.

A signal’s informational content can also vary in its complexity and structure. Adapting an existing repertoire in new ways, such as creating vocal sequences, may be a more parsimonious solution to increasing the efficiency of information transfer than producing new vocalizations (Nowak and Krakauer, 1999). This may be particularly important for group-living species that experience a wider range of social contexts, thereby increasing the need to be able to transmit more complex information (Freeberg et al., 2012). There has been interest in group-living primates as model species for such research, as they partake in complex cooperative behaviors that require communication between the parties involved (Engesser and Townsend, 2019; Zuberbühler, 2018). An increasing number of studies suggest that animal communication systems possess qualities reminiscent of syntactic compositionality and combinatoriality (Bolhuis et al., 2018; Townsend et al., 2018). Researchers have focused on understanding communicative system properties reminiscent of those of human language due to the inherent interest in understanding its origin and evolutionary path. While this area of study has been fruitful, the approach remains controversial as combinatorial organization is far more limited in comparison to the capabilities of human language, creating debate pertaining to the manner in which to interpret these findings (Scott-Philips & Heintz, 2023). The pursuit of understanding this intricate area of comparative linguistics may also divert attention away from the large variation of simpler nonhuman animal vocalizations. Another, more simple vocalization organization, that might act as a precursor to complex communication, may be the call sequence. Sequences can be composed of a number of distinct element types or of repeats of the same element and the information stored in the sequence may or may not reflect the complexity of its structure (Kershenbaum et al., 2016). The elements composing a given sequence could potentially hold different informational content and, as a result, play different functional roles.

Common marmosets (*Callithrix jacchus*) are monomorphic primates. They are cooperative breeders living in large social groups, that display a wide range of complex social behaviors, such as allomaternal care, group vigilance and active food sharing (Burkart et al., 2022; Digby et al., 2010; Saito, 2015). Common marmosets possess a rich vocal repertoire that help regulate these intricate interactions (Agamaite et al., 2015; Bezerra and Souto, 2008). They not only emit basic tonal signals but also produce intricate calls composed of multiple elements (Bakker et al. 2018) and have the capacity to interconnect these sounds to create call combinations (Burkart et al. 2018; Bezerra & Souto, 2008; Cleveland & Snowdon 1982). Furthermore, it has been demonstrated that they display vocal flexibility through alterations in call duration (Brumm et al., 2004), intensity (Brumm et al., 2004; Eliades & Wang, 2012, but see Pomberger et al., 2020), complexity (Pomberger et al., 2018), and timing (Pomberger et al., 2018; Roy et al., 2011). Finally, they regularly produce sequences of calls. Marmoset phees (long-distance contact calls) can be produced singularly as well as in a sequence (repeats of just the phee call type or in combination with other call types). Phees have been the subject of a wide range of studies (e.g., An et al., 2023; Chen et al., 2009; Miller et al., 2010; Miller and Wren, 2012; Norcross and Newman, 1993; Yamaguchi et al., 2010; Zürcher et al., 2021), however, these studies widely regard phee elements as equivalent and have not considered them in the context of sequences, their co-occurrence within one bout or the order in which they are emitted.

The aim of this project was to investigate whether the structure and informational content of common marmoset phees change depending on their position within a sequence. We tested the consistency of conveying identity and sex information across elements in a sequence. Research into acoustic sequences contributes to the growing body of evidence of the diversity of complex structures in animal communication systems and may help elucidate the evolution of human language.

## Methods

### Study subjects

We recorded phee calls (long-distance contact call produced when group members have been separated from one another (see Agamaite et al., 2015 for a description of the call’s characteristics) from 6 adult common marmosets (3 male and 3 female) housed at the University of Zürich between January and August 2022. Throughout the day animals were supplied with water *ad libitum*. They were fed vitamin-enriched mash and/or fresh fruits and vegetables at least twice a day, once in the morning, and once around noon. In the afternoon they received additional protein-rich food morsels in the form of either insects, gum, boiled eggs, or cheese. The subjects were part of groups or pairs that were housed in separate heated indoor enclosures (2.4x1.8x3.6m) and had access to corresponding outdoor enclosures of the same dimensions. All procedures were done in accordance with Swiss legislation and were licensed by Zürich’s cantonal veterinary office (license ZH223/16).

### Recording procedure

Individuals were recorded two at a time (always in a male and female combination), when placed in adjacent wire cages. All male-female combinations (6 individuals, 9 combinations) were recorded 12 times spread across 8 months, with each recording lasting 15 minutes. We used two CM16/CMPA - Avisoft Bioacoustics condenser microphones for all audio recordings and annotated caller identity in real-time using Avisoft Recorder (Specht, 2002). We could visually distinguish between the different study subjects through distinctive markings on their fur. We were able to record their identity in real time through the Avisoft Recorder.

### Data processing

We used Avisoft-SASLab Pro *ver. 5*.*3*.*01* (Specht, 2002) for call extraction. We extracted phees occurring alone (type A; Figure 1A), as well as in sequences (type B and C; Figure 1B and 1C). Sequences were defined as a series of elements occurring within 0.5s. A sequence always contained at least one phee element but could also include other call types surrounding or separating the main phee elements. If another element (either phee or other call type) occurred within a 1s interval before the start or beginning of a sequence, and there were no other calls present for a given period before or after it (we set this to equal the duration of the whole vocalization bout) we also included it as part of the sequence. We noted the identity of the caller and the numerical position of the phee element within its respective sequence. For analysis, we retained single phees and those occurring in two- or in three-phee-element sequences as higher number sequences occurred infrequently. Only vocalizations which could reliably be assigned to a given caller were kept. We also excluded low-quality noisy calls that overlapped with each other or with interfering background noise (e.g., cage rattling).

**Figure 1:**
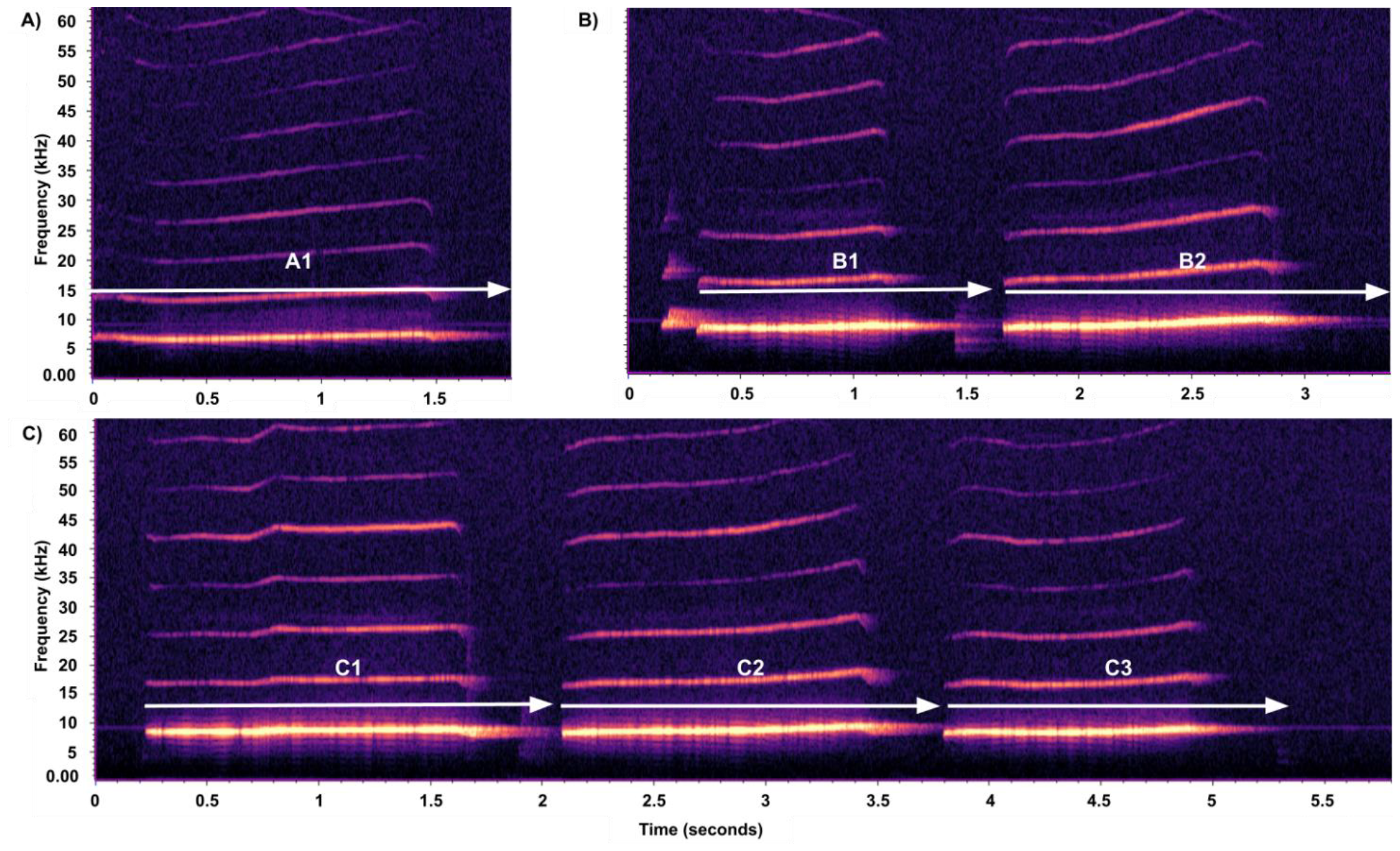
Examples of spectrograms of A) a single phee; B) a B-type sequence; C) a C-type sequence. Labels indicate which sequence type the element originated from (A, B or C), and its relative position within the sequence (e.g., single phees = A1, elements in the second position of a B-type sequence = B2, etc.). Spectrogram settings: Hann window size of 400 ms, hop size of 200 ms, and overlap 50%.

For acoustic parameter extraction, we used Raven Pro *ver. 1*.*6*.*4* (K. Lisa Yang Center for Conservation Bioacoustics; spectrogram settings: Hann window size of 400 ms, with a hop size of 200 ms, and overlap 50%). We used a bandpass filter (between 2kHz and 10kHz) to ensure the extraction of data based on a call’s fundamental frequency and to diminish noise that could affect the accuracy of the measurements. We used robust measurements (Table 1, Figure 2) as they are less sensitive to observer bias and the exact position of the selection box boundaries, by relying on the energy stored within the selection (Charif et al., 2010; Cortopassi, 2006).

**Table 1:** Acoustic parameters generated by Raven Pro used in analyses. Definitions were taken from Charif et al. (2010).

| Parameter (units) | Definition |
| --- | --- |
| **Average Entropy (bits)** | Average Entropy is calculated by finding the entropy for each frame in the selection and then taking the average of these values. Higher entropy values correspond to greater disorder in the sound whereas a pure tone with energy in only one frequency bin would have zero impurity. The Average Entropy measurement describes the amount of disorder for a typical spectrum within the selection. |
| **Frequency 5% (Hz)** | The frequency that divides the selection into two frequency intervals containing 5% and 95% of the energy in the selection. |
| **Frequency 95% (Hz)** | The frequency that divides the selection into two frequency intervals containing 95% and 5% of the energy of the selection. |
| **Center Frequency (Hz)** | The frequency that divides the selection into two frequency intervals of equal energy. It can be stated that it is the smallest discrete frequency in which the left side of the formula exceeds 50% of the total energy in the selection. |
| **Bandwidth 90% (Hz)** | The difference between the 5% and the 95% frequencies. |
| **Duration 90% (s)** | The difference between the 5% and the 95% times. The 5% Time and 95% Time are the points in time which divide the selection into two time intervals containing 5% and 95% and 95% and 5% of the energy, respectively. |
| **Center Time (s)** | The point in time at which the selection is divided into two time intervals of equal energy. It can also be stated that it is the smallest discrete time in which the left side of the formula exceeds 50% of the total energy in the selection. |

**Figure 2:**
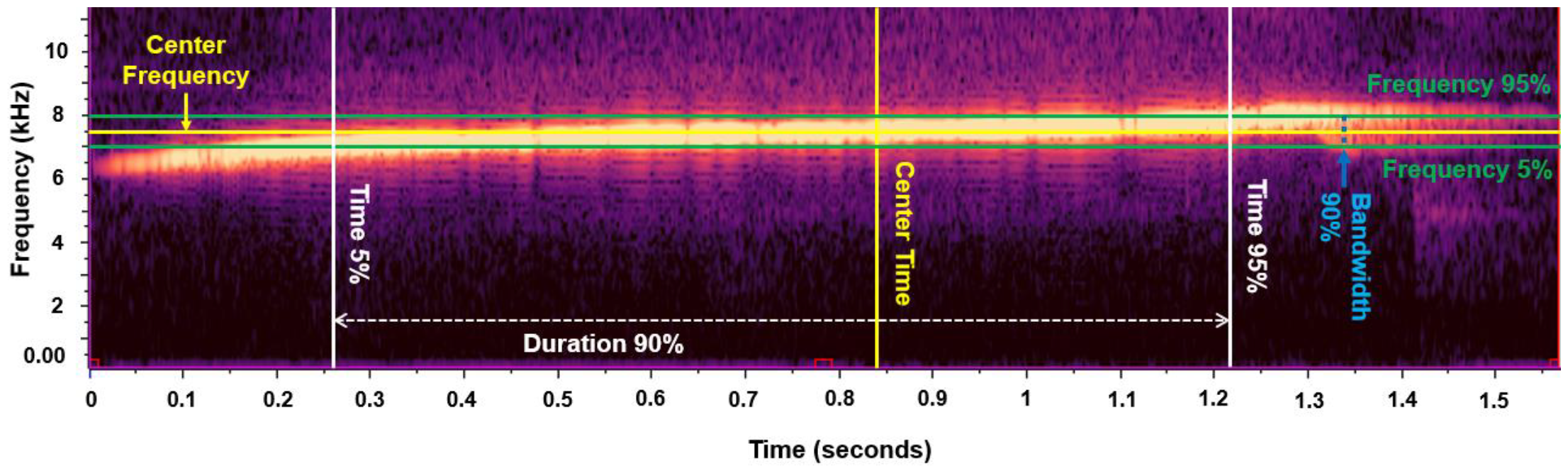
Spectrogram visualization of the spectral-temporal parameters used in the analysis. Spectrogram settings: Hann window size of 400 ms, hop size of 200 ms, and overlap 50%.

To the list of parameters, we also added a variable describing the “shape” of the phee, as seen on a spectrogram (i.e., whether the signal was linear or curved; if it was curved, whether it followed a convex or concave trajectory; and whether the slope was positive or negative). To do this, we extracted the values laying on the 10^th^ to 90^th^ percentiles along the length of the Peak Frequency Contour (a list of Peak Frequency estimates across the length of a selected call; Charif et al., 2010). Then, we created a linear regression model with a quadratic term to describe the nature of the relationship between these values. We interpreted a non-significant linear model term as a flat line. If the term was significant, we used the sign of the slope of the line as an indicator of directionality. A significant quadratic term indicated the presence of a curve, and we used the sign of the slope of the quadratic term to determine whether the curve was convex or concave.

### Data analysis

All data were analyzed in R (*ver. 4*.*2*.*3*). To examine the informational content and the presence of structural differences between different phee element types, we created supervised random forest classifiers using the *randomForest* and *caret* R packages (Kuhn, 2008; Liaw and Wiener, 2002). We chose this approach, as it has been shown to be robust to unbalanced and noisy datasets, can be used to identify the features that contribute most to accurate classification in the model, and can utilize large non-linear and multidimensional data sets (Breiman, 2001; Valletta et al., 2017).

To evaluate each model’s performance, we split the data into training (70% of samples) and testing (30% of samples) subsets according to the classifier being tested (i.e., if using all available phees to classify to identity, the training subset contained 70% of each individual’s total calls). For each model, we independently tuned the *mtry* hyperparameter to ascertain the optimal number of variables used at each split that would improve overall performance, and otherwise used the default settings. Additionally, the *randomForest* package provides estimates for the importance of each of the features, by determining how much prediction error would increase if that variable were changed (Liam and Wiener, 2002). For the purpose of this study, we set the threshold for accuracy to 70% to consider the machine accurate in classifying a datapoint to a given class.

To determine whether there are any structural differences between elements according to their relative position within and between sequence types we created separate random forest classifiers to test whether it was possible to accurately distinguish between: (1) B1 and B2 elements; (2) C1, C1, and C3 elements; (3) single A1 elements, B1, and C1 elements; (4) B2 and C2 elements. The datasets used in these models were unbalanced in terms of number of calls per individual and per sex. To deal with the issue of non-independence in the data, we also added identity and sex as variables. If the results from the classifiers suggested we could reliably distinguish between elements from the same sequence type then we constructed a linear mixed model (LMM) with the most important model feature that was a spectral-temporal parameter. This would help explain what drives the observed difference between elements. We constructed two such models, one examining when Center time (s) occurs in B1 and B2 elements, and a similar test for C1, C2, and C3 elements for their square root values of Center time (s).

We determined the robustness of identity and sex information by using two separate random forest models to classify all phees in our data set to a given individual or sex. Finally, we determined whether the accuracy of classification of phees to a given caller or sex varied across different element types (e.g., whether it varied for A1, B1, B2, C1, C2, and C3 elements). We did this by creating separate classifier models for each of these element types (for both the case where we classify a phee to an individual and to a sex) and comparing their accuracy scores. For each of the classifier models we subsetted the data for the relevant element type but otherwise ran the same model. In the models classifying to identity, we input sex as an additional feature, to account for possible non-independence. We decided to exclude one female individual from all models, due to the low sample size of calls produced by this individual.

## Results

The final dataset we used to run the models consisted of 3066 phees (1525 and 1541 calls produced by females and males, respectively), with 613 calls per individual, on average (range: 446-781 calls).

We were able to classify phees to a sex with an accuracy of 88.49% and OOB error rate of 11.66% (n = 2230). The most important model feature for classifying to sex across all phees was Frequency 5% (Hz). Phees could be classified to an individual with an accuracy of 83.15% and OOB error rate of 16.03% (n = 3066). The most important feature in the model classifying all phees to identity was sex of the caller.

The random forest algorithm classified first and second position elements in B-type sequences into two separate clusters with an accuracy of 83.42% and OOB error rate of 13.15% (n = 1347). The most important source of variation in the model was the Center time feature. Additionally, the LMM showed that Center time occurred later in first elements than in second (LMM, F = 785.09, p < 0.0001; Figure 3A). The models could cluster the elements within C-type sequences with an accuracy of 76% and OOB error rate of 24.27% (n = 1414). Center time remained the most important feature. More specifically, the square root value of Center time occurs later in first than in second (Tukey test, p < 0.0001) or third elements (Tukey test, p < 0.0001) and later in second than in third elements (Tukey test, p < 0.0001; Figure 3B).

**Figure 3:**
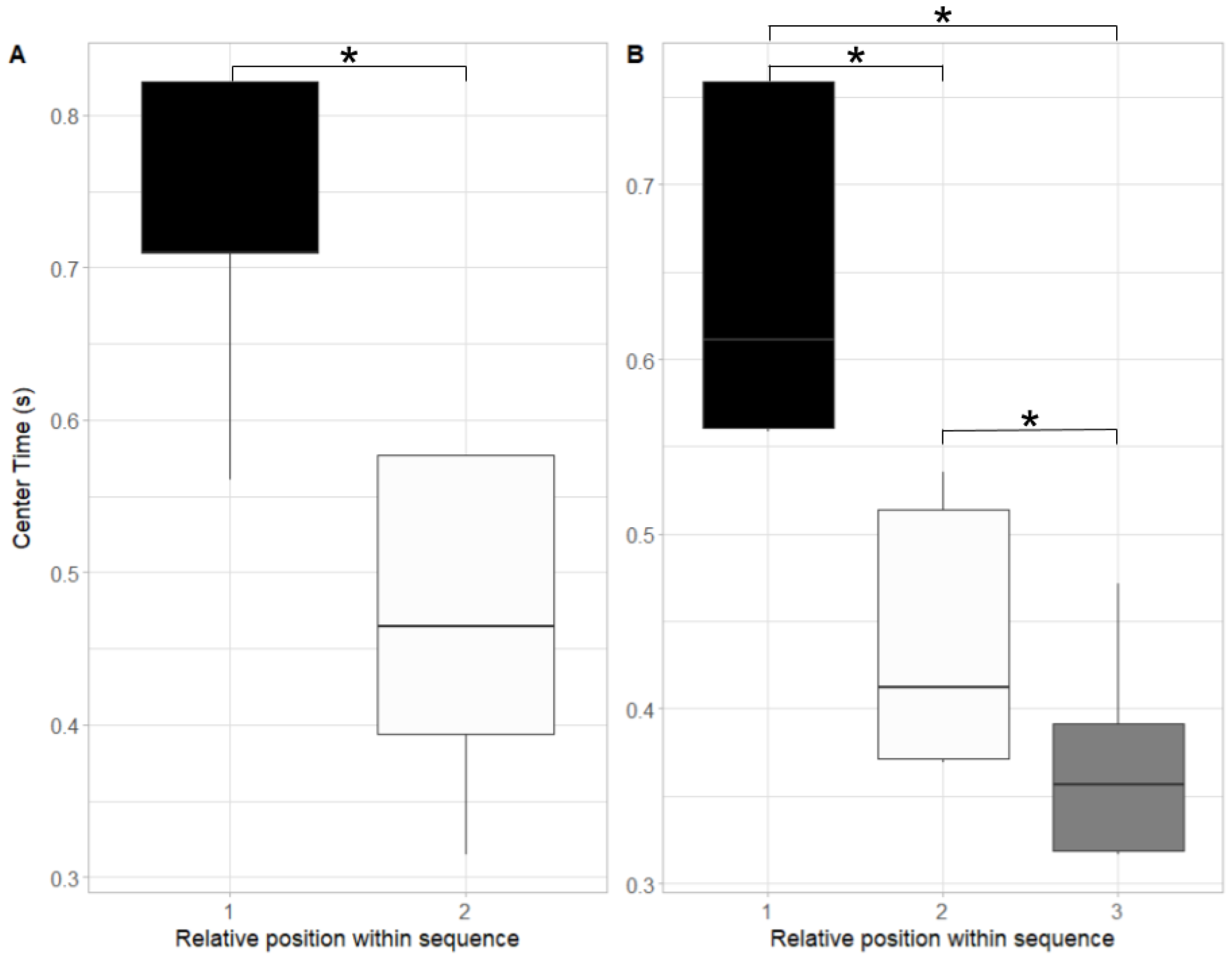
Center time (s) difference between A) first and second position elements in two-element phee sequences; B) first, second, and third position elements in three-element phee sequences. Predicted model values for Center time are plotted. Asterisks indicate significant pairwise comparisons.

The first elements of sequences of different lengths were similar to each other (model accuracy = 60.38% and OOB error rate = 36.93%; n = 1415). The model can classify second elements to B- and C-type sequences with an accuracy of 73.65% and OOB error rate of 24.43% (n = 1175). The most important feature in both models was identity. The results from the models classifying identity and sex, respectively, for each element type are presented in Table 2. We observed higher identity accuracy rates in second elements and higher sex accuracy rates in first elements.

**Table 2:**
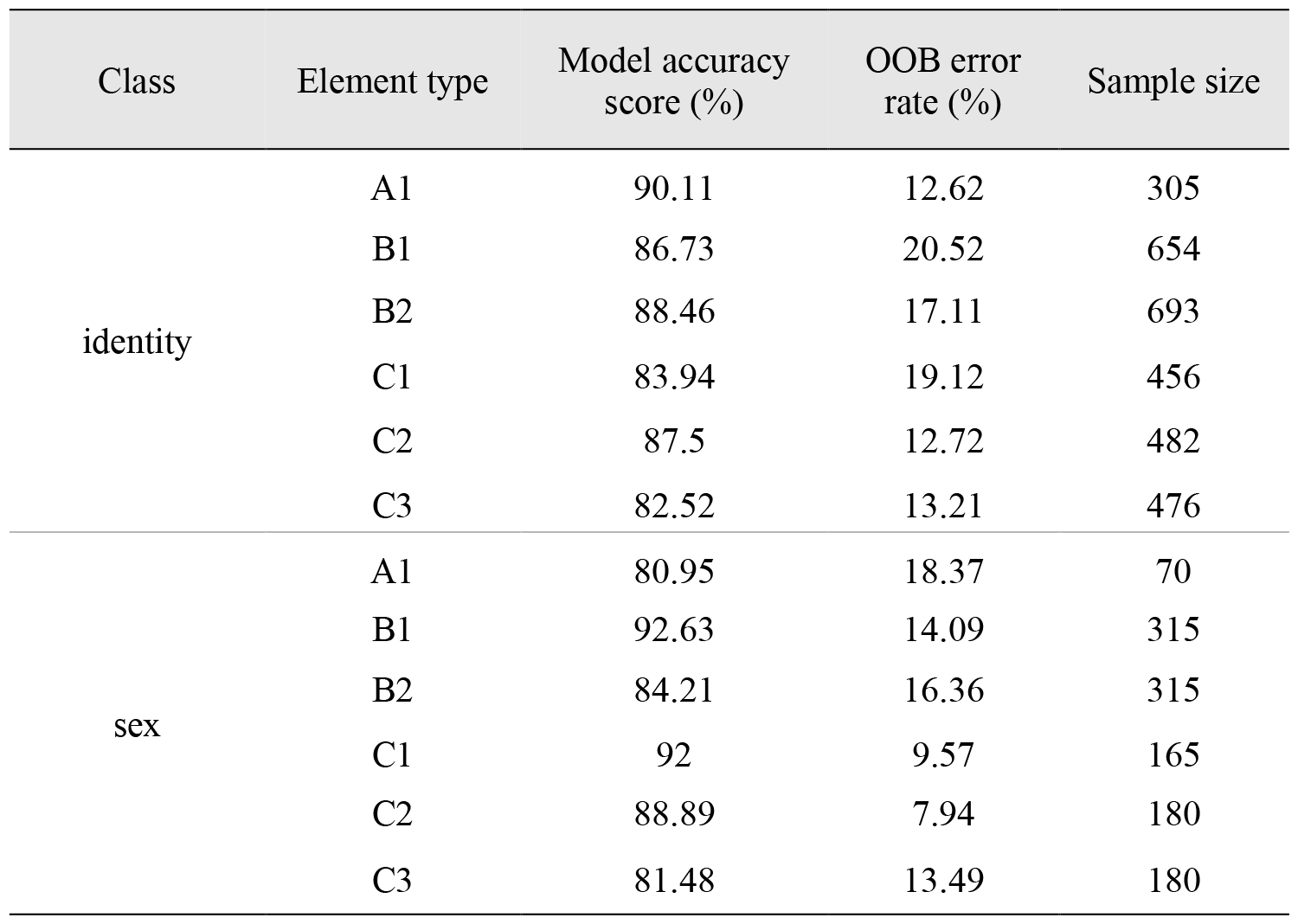
Accuracy of phee element type classification to caller identity and sex.

| Class | Element type | Model accuracy score (%) | OOB error rate (%) | Sample size |
| --- | --- | --- | --- | --- |
| identity | A1 | 90.11 | 12.62 | 305 |
|  | B1 | 86.73 | 20.52 | 654 |
|  | B2 | 88.46 | 17.11 | 693 |
|  | C1 | 83.94 | 19.12 | 456 |
|  | C2 | 87.5 | 12.72 | 482 |
|  | C3 | 82.52 | 13.21 | 476 |
| sex | A1 | 80.95 | 18.37 | 70 |
|  | B1 | 92.63 | 14.09 | 315 |
|  | B2 | 84.21 | 16.36 | 315 |
|  | C1 | 92 | 9.57 | 165 |
|  | C2 | 88.89 | 7.94 | 180 |
|  | C3 | 81.48 | 13.49 | 180 |

## Discussion

Here we showed that all common marmoset phee elements encode information about the caller’s sex and identity, however the amount of this information encoded varies between elements within the same sequence. Our results suggest that first phees of multi-elements sequences might be used by the receiver to narrow down the list of potential callers (based on sex information), which can later be more accurately identified through the subsequent phee elements. We showed that within- and between-sequence type differences between call elements exist, with the exception of the first elements of each sequence, which are similar in terms of their acoustic structure across sequences of different lengths. In this study we demonstrated that single phees and sequences consisting of phee elements are structurally and informationally diverse despite being comprised of the same call type. Our result provide evidence that repeat sequences might be functionally complex structures and provide an interesting subject for future research into animal communication systems and language evolution.

The random forest algorithm differentiated calls produced by males and females. For many organisms the ability to differentiate between sexes may be useful during intrasexual competition and mate attraction. Female-female interactions may be agonistic in socially monogamous cooperative breeders, such as the common marmosets, because of resource limitation and high reproductive costs (Clutton-Brock et al., 2006; Saltzman et al., 2009; Yamamoto, 2005). Subordinate females do not inherit the breeding position and are usually expelled from their natal group (De Sousa et al., 2009). To avoid aggressive encounters, it may benefit dominant females to monitor for immigrant competitors in their vicinity (Bezerra et al., 2007; Lazaro-Perea et al., 2000). On the other hand, sex signatures may allow outsiders to keep track of group composition, aiding them in detecting the opening of a breeding vacancy (Lazaro-Perea et al., 2000; McComb and Reby, 2005). Signaling sex may also benefit recently emigrated individuals in identifying who to form pair-bonds with and may help inform already pair-bonded individuals of the presence of potential mates to engage in extra-pair copulation with (Miller et al., 2004; Rubow et al., 2017).

In mammals, differences in fundamental frequency between sexes has often been attributed to body size differences (Charlton et al., 2020; Rendall et al., 2004). However, there is also evidence for such variation in monomorphic species, which may be explained by differences in vocal apparatus morphology (Frydman et al., 2023; Lenell and Johnson, 2017). Previous work investigating sex signature presence in phees has produced ambiguous results (Miller et al., 2010; Norcross et al., 1999; Norcross and Newman, 1993). Norcross et al., (1993) suggested that the sexes can be distinguished through frequency differences. Our results at first corroborate this, as Frequency 5% (or in practical terms, the fundamental frequency; Wierucka et al., 2021) is the most important feature. However, the LMM fails to detect frequency differences between the sexes. The random forest model is informed by 7 spectral-temporal features, whereas the LMM used only one feature. It is possible that the variation explained solely through the fundamental frequency is too small to be picked up by the LMM, and that the sex signature is dependent on an array of spectral-temporal features. As outlined in the previous paragraph, the ability to signal sex is important in regulating various social interactions. If this signature is encoded through multiple spectral-temporal features, instead of just one (i.e., the fundamental frequency), this may make it more robust to transmission failure and loss of information, thus ensuring that the signal is successfully conveyed to the receiver(s).

Consistent with earlier acoustic analyses investigating phees (Miller et al., 2010; Norcross, 1993), we could accurately predict the individual identity of the caller. Individual recognition underlies a myriad of social mechanisms, and it is not surprising that we find it here. It has been widely investigated in a variety of model organisms, including different primate taxa, such as chimpanzees, rhesus macaques, baboons, and other callitrichid species, which supports the idea that it is a fundamental component of complex sociality and group living (Kondo and Watanabe, 2009). Phees can be produced as territory advertisement, and during antiphonal exchanges which occur when individuals have been separated and need to remain in contact with the group (Agamaite et al., 2015; Bezerra and Souto, 2008). Common marmosets display an array of complex social behaviors which require collective cohesion and coordination between group members (Burkart et al., 2022; Digby et al., 2010). Individual recognition enables individuals to monitor the contributions of other groupmates and to alter their own behavior in response (Sharpe et al., 2013). We also show that all elements within a sequence encode identity which may ensure that this information is successfully conveyed to the receivers in the event of group separation and other instances of signal obstruction.

The most important feature used in the models classifying to identity was sex. This is consistent with previous results obtained by Phaniraj et al., (2022). Machine learning algorithms were shown to perform better when following a hierarchical approach to classification by first clustering by categories with higher accuracy, followed by less reliable classifiers. Our results suggest that information about the sex of a source caller is conveyed more accurately than identity. It follows that the random forest model would place high importance on this feature. The spectral-temporal features used to inform the model classifying to identity across all phees appear to hold similar weight. This could imply that interindividual differences are not determined through variation in a single parameter, in line with what has been previously suggested for male African savanna elephant social rumbles (Wierucka et al., 2021).

Our results indicate that the differences between elements within the same sequence are more prominent than those between elements in the same relative order in different sequences suggesting these elements do not hold equivalent functional roles. Elements in the first position of different sequence types are more structurally similar to each other, than are second position elements. Additionally, the accuracy of static information signatures (identity and sex) varies depending on which element type we examine. Sex information is more encoded than identity in first position elements from multi-element sequence types (i.e., B1 and C1 elements). In contrast, the identity scores of second position elements (i.e., B2 and C2) are higher than first position elements. Consequently, it seems that the emphasis on the information transmitted is first on the sex of the caller, followed by their identity.

Sex is more strongly encoded than identity and models classifying to identity rely most on the sex feature for accurate classification (Phaniraj et al., 2022). From a functional standpoint, it may make sense for there to be a sequential order for vocal signature recognition. A previous study done on Australian sea lion mother-pup pairs showed that mothers were able to distinguish between pups of different age classes based on visual cues, before identifying them individually through acoustic or olfactory cues (Wierucka et al. 2018). These findings show a reliance of sea lion females on first refining the recognition process to correctly identify the broader group that the individual they are searching for belongs to. Likewise, common marmosets may first classify an individual to a wider group denominator through the information encoded in the first elements from multi-element sequences, which convey more strongly the sex of the caller. As a result, the receivers may be narrowing down the options for recognition to a subset of conspecifics, thereby refining the individual identity recognition process and making it more robust against potential error.

Our results suggest that identity is more accurately encoded than sex in single phees, despite A1, B1, and C1 elements showing structural similarity (which may have led to the expectations that their informational content might also be the same). As these phees are produced singly with no other elements to complement their informative function and aid in narrowing down the recognition process it follows that accurately conveying information about caller identity is their most important function (Miller et al., 2010; Norcross and Newman, 1993; Zürcher et al., 2021). While we did not investigate if and how additional vocal signatures pertaining to a caller’s characteristics such as age, group identity, quality, etc., are encoded in single phees and in sequences it would be interesting to examine how they may change the pattern we observed here. Determining whether additional information about the caller, outside of their sex and identity, influences the recognition process, and if so – how exactly this is achieved, was beyond the scope of this study but would be an interesting topic for future work.

Generally, same-element sequences are thought to convey different information depending on changes in the rate of repeats or in the length of inter-element intervals (Engesser et al., 2017; Engesser and Townsend, 2019; Manser, 2001; Zuberbühler, 2018). Phee sequences do not seem to cleanly fit into this classification: 1) single phees are meaning-bearing units, as demonstrated by previous work and this project; 2) each element within a sequence displays variation in acoustic structure and signature accuracy; 3) sequences follow consistent inter-element interval patterns (Huang et al., 2022); 4) so far, there is no evidence that the function of a sequence changes according to the number of repeats (Engesser and Townsend, 2019; Kershenbaum et al., 2016), making them less similar to classical repeat sequences. The findings here may warrant more thorough research into the functions of repeat sequences and their constituent elements outside of their most studied alert function associated with increased state of arousal (Kern and Radford, 2013; Kershenbaum et al., 2016; Wheatcroft, 2015) in other taxa. Researchers in the field have not paid much attention to repeat sequences most probably because they do not represent a conventional analogue to the combinatorial properties of human language and hence might not be appropriate for the study of its evolution. However, our results and other recent research on pied babbler ‘clucks’ and ‘purrs’ by Engesser et al. (2017) reveal potential variation in the functionality of repeat sequences thus contributing to the diversity and complexity of animal communication systems. One way to facilitate cooperative behaviors is through vocal communication between individuals. As the field of animal communication grows, so does the body of evidence that compositionality and combinatoriality are widespread abilities across different animal taxa and may not just be properties of human language (Engesser and Townsend, 2019; Kershenbaum et al., 2016). Common marmosets form part of complex social networks which require frequent interactions and collective action between group members (Burkart et al., 2022). The findings in this project may possibly provide evidence for syntactic-like structural abilities by showing that the constituent elements of phee sequences do not hold equivalent structural or informative value. Studying the diversity of animal vocal abilities and their function in social organization may help us elucidate their selective drivers and understand human language’s evolutionary history.

## Acknowledgements

Funding: KW & JMB were partially or entirely funded by the NCCR Evolving Language, Swiss National Science Foundation Agreement #51NF40_180888.

